# Benchmarking Imputed Low Coverage Genomes in a Human Population Genetics Context

**DOI:** 10.1101/2024.06.02.597067

**Authors:** Gludhug A. Purnomo, João C. Teixeira, Herawati Sudoyo, Bastien Llamas, Raymond Tobler

## Abstract

Ongoing advances in population genomic methodologies have recently made it possible to study millions of loci across hundreds of genomes at a relatively low cost, by leveraging a combination of low-coverage shotgun sequencing and innovative genotype imputation methods. This approach has the potential to provide economical access to genotype information that is similar to most widely used low-cost genotyping approach – i.e. SNP panels – while avoiding potential issues related to loci being ascertained in distantly related populations. Nonetheless, adoption of imputation methods has been constrained by the lack of suitable reference panels of phased genomes, as performance degrades when panel individuals are distantly related to the target populations. Recent advances in imputation algorithms now allow genetic information from the target population to be used in the imputation process, however, potentially mitigating the lack of a suitable reference panel. Here we assess the performance of the recently released GLIMPSE imputation software on a set of 250 low coverage genomes (∼3x) from populations from Island Southeast Asia and Near Oceania that are poorly represented in publicly available datasets, comparing the use of imputed genotypes against other common genotype calling methods for a range of standard population genomic analyses. We find that imputation performance and inference both greatly improved when genetic information from the 250 target individuals was leveraged, with comparable results to pseudo-haploid calls that trade off improved precision with reduced accuracy. Our study shows that imputed genotypes are a cost effective and robust basis for population genomic studies of groups, especially those that are poorly represented in publicly available data.

## INTRODUCTION

Population genomic studies of human evolution and demographic history have continued to expand in geographic coverage over the last decade, including the first genomic surveys of populations in Island SouthEast Asia (ISEA) and Near Oceania (i.e. New Guinea, Australia and Island Melanesia) (Friedlaender et al., 2007; Hill et al., 2007; Ingman & Gyllensten, 2003; van Holst Pellekaan et al., 2006). Because of high costs associated with generating deeply sequenced genomes (i.e. ≥30x coverage) on a population scale, only a handful of population genomic studies of ISEA and Near Oceania have utilized deep coverage whole genome sequencing (Brucato et al., 2021; GenomeAsia100K Consortium, 2019; Jacobs et al., 2019; Malaspinas et al., 2016). Instead, many population genomic studies in this region have leveraged mitochondrial DNA (Delfin et al., 2014; Gomes et al., 2015; Pedro et al., 2020; Tumonggor et al., 2013) and single nucleotide polymorphism (SNPs) arrays to capture population-level genomic information (Hudjashov et al., 2017; HUGO Pan-Asian SNP Consortium et al., 2009; Kusuma et al., 2017; Larena et al., 2021), often augmenting the SNP set via imputation from a genomic reference panel of worldwide human diversity (Byrska-Bishop et al., 2021; Delaneau et al., 2014). While being a cost-effective means of generating population genomic data, the effectiveness of this approach is hindered by usage of SNP arrays that use loci ascertained in populations comprising predominantly European ancestry (these SNPs are often less informative about ISEA/Near Oceania population history) and genomic reference panels that lack representation of ISEA and Near Oceania (reducing the precision of imputation) (Lachance & Tishkoff, 2013).

In the past decade, the emergence of imputation methods that work with low coverage whole genome sequencing (e.g. Beagle (Browning et al., 2018), Minimac2 (Fuchsberger et al., 2015), Impute2 (B. Howie et al., 2012), Glimpse (Rubinacci et al., 2021)) has seen this approach being proposed as a viable alternative to imputed SNP arrays as a cost effective means of capturing population genomic diversity that may overcome some of the drawbacks associated with the latter (Browning & Browning, 2016; Fuchsberger et al., 2015; B. N. Howie et al., 2009; Rubinacci et al., 2021). Low coverage WGS (henceforth, lc-WGS) has been repeatedly shown to outperform standard SNP arrays when combined with genotype imputation in the context of genome-wide association studies (GWAS), performing at least as well for common variants and providing significantly better coverage of rare variants (CONVERGE consortium, 2015; Gilly et al., 2016). Moreover, in the past few years new imputation methods have emerged that allow genetic data from target populations to be used in the imputation process (e.g. BEAGLE (Browning & Browning, 2016), STITCH (Davies et al., 2016), GLIMPSE (Rubinacci et al., 2021). Crucially, this latter feature means that imputation can still achieve high accuracy even when relevant population information is missing from available reference panels (Byrska-Bishop et al., 2021; Delaneau et al., 2014).

The combination of low coverage WGS and robust imputation methods holds the promise of greatly expanding the scope and scale of population genomic research without compromising statistical power or inflating artifactual results. However, while these properties have been confirmed in the GWAS context, systematic exploration of the efficacy of low coverage WGS imputation methods for population genomic inferences of demographic and evolutionary history are still rare (e.g. (Lou et al., 2021)). In this study, we benchmark the precision and performance of imputed genotypes from low coverage sequence data (∼1-4x) from >250 genomes from populations ISEA and Near Oceanians for three widely used population genomic inference procedures – i.e. PCA (Patterson et al., 2006; Price et al., 2006), ADMIXTURE (Alexander & Lange, 2011), and *f*4 statistics (Patterson et al., 2012; Peter, 2016) – and compare this to the performance of ‘naive’ genotypes (i.e. most probable genotype call; see Methods) and also pseudohaploid calls (i.e. random sampling of a single allele at a given locus; see Methods) that are standard in the ancient DNA field where low coverage sequence data is the norm. Imputation is performed using the GLIMPSE algorithm, which is one of the best performing imputation methods designed to work with low coverage WGS data (Rubinacci et al., 2021), while also drawing upon genetic information from target samples during genotype imputation, which is crucial for ISEA and Near Oceanian populations whose ancestry is poorly represented in available reference panels. We determine the best performing genotyping approaches through comparison of genotype calls from low coverage genomes to a truth set obtained from high coverage sequencing (see **Materials and Methods**) and discuss the broader implications of low coverage genotyping for inferring human population genomic history and evolution.

## MATERIAL AND METHODS

### Sample collection and ethics

The genetic data for this project comes from 256 individuals from 11 different populations across Wallacea and (i.e., Kei, *n* = 20; Aru, *n* = 23; Tanimbar, *n* = 22; Seram, *n* = 27; Ternate, *n* = 30; Sanana, *n* = 19; Daa [in Central Sulawesi], *n* = 22; and Rote-Ndao, n = 29) and West Papua (Keerom, *n* = 26; Mappi, *n* = 11; and Sorong, *n* = 27). Permission to conduct the research was granted by the National Agency for Research and Innovation, under the auspices of the Indonesian State Ministry of Research and Technology. Informed consent for all 256 individuals was obtained for the collection and use of all biological samples during community visits that were overseen by the Indonesian Genome Diversity Project (IGDP) team, following the Protection of Human Subjects protocol established by Eijkman Institute Research Ethics Commission (EIREC). The study is also approved by The University of Adelaide Human Research Ethics Committee (Ethics approval no. H-2020-211).

### Whole genome sequencing, read processing and alignment

DNA was extracted from whole blood samples for all 256 samples at the Eijkman Institute for Molecular Biology Jakarta using the Gentra Puregene Blood Core Kit C (QIAGEN) following the manufacturer’s protocol. DNA sequence libraries were prepared using the Nextera DNA Flex Library Preparation Kit (Illumina) following the recommended protocol. After quantifying DNA concentrations for each sample using Qubit dsDNA BR Assay Kit (Thermo Fisher Scientific), all 256 samples were multiplexed into a single pool and submitted to 150 bp paired-end sequencing across three lanes of Illumina NovaSeq S4 flowcell. To obtain high (∼30x) coverage samples for comparison, eight of the 256 samples were chosen for sequencing on a single NovaSeq S4 lane, with one sample coming from each of the following populations (sample ID in parentheses along with specific island/region of origin if this is not the same as the population label) – Seram (SRM-HLU013); Daa (Sulawesi; KAL007); Rote (RTE045); Sanana (TNT160); North Maluku (TNT172); Aru Island (ARU-LRK007); Sorong (West Papua; SRG059); and Keerom (West Papua; KRM048).

For all samples sequenced at high coverage, raw sequence reads in the fastq format were pre-processed with fastp to remove adapters and trim poly-G and poly-X tails, with first 20 and last 5 nucleotides being trimmed if they fell below a quality threshold of 20 (Chen et al., 2018). The sequencing reads were then mapped and processed following the recently published protocol of the Human Genome Diversity Panel (HGDP) dataset, as outlined in Bergstrom et al 2020. Briefly, processed reads were mapped to the human reference genome GRCh 38 (hg38) using BWA mem v0.7.17 with the -T 0 parameter (Li, 2013). Mapped sequencing reads were sorted and duplicated reads marked using biobambam2 (Tischler & Leonard, 2014), with nucleotide bases recalibrated using baseRecalibrator from the GATK software suite v3.5 (McKenna et al., 2010).

The low coverage genomes were pre-processed and mapped using the same protocols outlined above, except all samples were merged into a single file using samtools v1.9 (Li et al., 2009) prior to the sorting and duplicate marking step of the merged reads.

### Determining ‘truth’ genotype set using high coverage sequencing data

The comparisons require determining a set of high quality genotypes for each of the eight samples, which are used for benchmarking the performance of different types of genotype calls in low coverage data. To obtain a set of high quality ‘truth’ genotypes, we replicated the relevant protocols used for the recently published HGDP dataset, as outlined in (Bergström et al., 2020). Briefly, single nucleotide polymorphisms (SNPs) and indel variants were called using GATK Haplotypecaller and GenotypeGVCFs (Poplin et al., 2018). Genotypes were set to missing if the genotype quality (GQ) was equal to or lower than 20, or the coverage depth (DP) was equal to or greater than 1.65 times the genome-wide coverage for each sample.

Next, the GATK Variant Quality Score Recalibration tool (VSQR) was used to compute call quality annotations (QD, MQRankSum, ReadPosRankSum, FS, MQ, VQSRMODE) to SNPs (files: hapmap_3.3.hg38.vcf.gz, 1000G_omni2.5.hg38.vcf.gz, and 1000G_phase3.snps.highconfidence.hg38.vcf.gz) and indels (files: Mills_and_1000G_gold_standard.indels.hg38.vcf.gz and Homo_sapiens-assembly38.known_indels.vcf.gz) (all files available from: gatk.broadinstitute.org/hc/en-us/articles/360035890811-Resource-bundle). Regions with excess heterozygosity (ExcHet) were calculated on a per-allele basis using the bcftools fill-tags plugin (Danecek et al., 2021). All SNPs with a VQSR score below -8.3929, or indels with VQSR score below -1.0158, and ExcHet value of at least 60 (corresponding to a *p*-value of 10^-6^) were set to missing.

### Genotype calling in the low coverage data

For all low coverage samples, variant discovery was performed using Haplotypecaller with parameters “--ERC GVCF” and “--includeNonVariantSites” to ensure that monomorphic sites were retained, and intermediate gVCFs were generated for individual samples. GATK CombineGVCFs was used to amalgamate all gVCFs on a population basis, with joint variant calling on each amalgamated gVCF being performed using GenotypeGVCFs, thereby allowing variant information from all samples in each population to be used in individual genotype calls. This step produces VCF files with Phred-scale Likelihood (PL), a normalized form of genotype likelihood, that are used in the subsequent imputation process.

Four different types of genotype calls were generated for the eight low coverage samples that were compared against their high coverage counterparts. First, ‘naive’ genotype calls (denoted in results by ‘Genotypè label) were made by retaining SNPs with a genotype quality (RGQ and GQ) ≥ 20, with all other SNPs set as missing. For the second and the third set of genotype calls, GLIMPSEv1.1 (Rubinacci et al., 2021) was used to impute genotypes using the set of 3,202 phased genomes from the 1000 Genomes Project (TGP) sample collection maintained at the New York Genome Centre (Byrska-Bishop et al., 2021) as the reference panel (see next section). The key difference between the second and third set of genotypes is that imputation was either performed only using the eight samples that were also sequenced as high coverage (labeled as ‘Impute_8’) or by additionally including the complete set of 256 low coverage samples in the imputation process (labeled as ‘Impute_all’). The Impute_all strategy could potentially improve genotype call accuracy by leveraging genetic information from the large number of samples with sharing related ancestry. The fourth and final set of genotype calls were obtained by randomly sampling a single read at a set of predesignated variant positions observed in the high coverage samples (see Results), creating ‘pseudohaploid’ calls (labeled as ‘Pseudohap’). This is a standard practice in ancient DNA studies where endogenous DNA yields are low and is known to produce unbiased analyses at the cost of halving the potential information available at each locus (Green et al., 2010). Pseudohaploid calls were made using the sequenceTools software (https://github.com/stschiff/sequenceTools.git), which is widely used in paleogenomic research, with only reads having both base and mapping quality ≥30 being used in the random sampling process.

### Data imputation

Imputation was performed following the protocol outlined on the official GLIMPSE github repository website using default parameterisations (https://odelaneau.github.io/). First, genomic ‘chunks’ were defined by running the GLIMPSE_chunk algorithm on the 3,202 phased TGP genomes that were used as the imputation reference panel, with each chromosome being run independently to speed up computation. Genotype imputation was performed for each resulting chromosome chunk using GLIMPSE_phase. The resulting VCF files for each chunk were then merged together using Glimpse_ligate, with all imputed loci having genotype probabilities (GP) lower than 0.9 set to missing (using the BCFtools plugin).

### Genotype call comparison

Several different analyses were performed to evaluate the performance of the four different genotype calling methods (i.e., Genotype, Impute_8, and Impute_all, and Pseudohaploid). First, for the eight samples sequenced at low and high coverage, the four types of genotype calls from low coverage genomes were compared to the corresponding truth set calls obtained from the high coverage data. Performance was measured using two complementary statistical metrics, 1) precision, or the ratio of truth positive calls (TP) to the total number of called genotypes (which combines true and false positives (FP); i.e., TP / [FP + TP]) and 2) missingness, or the percentage of SNPs where genotypes could not be confidently called (i.e., where data was insufficient or filtered due to low quality). Comparisons were calculated separately for each genotype (i.e., homozygous reference , heterozygous , and homozygous alternative, reported here as ‘0/0’, ‘0/1’, and ‘1/1’, respectively) and also on the set of aggregated genotypes. Notably, it is not possible to obtain a meaningful comparison for heterozygous calls when using pseudohaploid data, since only a single allele is sampled. Instead, we calculated the degree of reference bias across all heterozygous SNPs for each sample, which captures the degree to which the reference allele is more likely to be sampled over the alternative allele due to an inherent alignment bias against alleles not present on the reference genome.

### Performance in population genomic analyses

Further comparative analyses were performed for three widely used population genetic methods: principal component analysis (PCA) using the smartpca function from EIGENSOFT v7.1.2 (Patterson et al., 2006; Price et al., 2006), ancestry estimation using ADMIXTURE v1.3.0 (Alexander & Lange, 2011), and *f*4 statistics using ADMIXTOOLS2 v2.0.0 package (Maier et al., 2022). For these analyses, the high coverage samples were merged with publicly available genomes from the Simons Genome Diversity Project (∼300 genomes from 142 different ethnic groups; SGDP (Mallick et al., 2016)), along with data from Indonesia (Jacobs et al., 2019), and New Guinea (Malaspinas et al., 2016), to create a comparative global dataset (Table S1). Because all genomes in this global dataset had been mapped to an earlier reference genome version (GRCh37 with decoy sequences), all SNPs were converted to GRCh38 coordinates using the liftover tool available in Genozip v.12.0.34 (https://genozip.readthedocs.io/dvcf.html; (Lan et al., 2022)) with the relevant chain file obtained from the UCSC Genome Browser. In this final merged dataset, SNPs missing in more than 5% of the combined samples, or having a minor allele frequency less than 1% across all samples, were removed using PLINK v.1.987 (Purcell et al., 2007), leaving a total of 5,166,352 SNPs available for further analysis. This merged dataset was further pruned to remove SNPs in moderate linkage disequilibrium (LD) with moderate levels of linkage (i.e., *r*^2^ > 0.4) on every 200 variant with 25 window shift were removed using PLINK v.1.987 (parameter indep-pairwise 200 25 0.4) (Lazaridis et al., 2016), which resulted in 528,617 SNPs. Both PCA and ADMIXTURE analyses were performed using the pruned SNP set, with *f*4 statistics measured on both the original and pruned SNP sets.

For the PCA, the first 10 principal components (PCs) were estimated for the combined set of global samples and eight high coverage genomes using the smartpca function with no outlier removal step (Hudjashov et al., 2017). Genotype calls from the eight low coverage samples were projected onto these 10 PCs. To test the optimal fit between low coverage genotypes and the truth set, Euclidean distances were measured between the low and high coverage genotypes for each sample (technically these are Mahalanobis distances, due to the transformation of the genotype space in the PCA).

For the ADMIXTURE analysis, ancestry components from *K* = 3 to *K* = 12 were estimated using the combined set of global samples and eight high coverage genomes using ADMIXTURE v1.30 (Alexander & Lange, 2011). Cross validation was used to estimate the optimal *K* coefficient, with eight ancestry clusters being found that best fit the genetic structuring in the global dataset (**Figure S1**). The inferences from the optimal ADMIXTURE run were subsequently used to estimate the representation of the eight ancestry components in the low coverage genotype data. The Euclidean distance between the values of the eight inferred ancestry components was measured between low coverage genotypes and the truth set in order to infer the optimal genotype calls for each sample.

Finally, *f*4 statistics of the form *f*(Truth, Target = Low coverage; Test = Papuan or East Asian, Africa) were computed using the ADMIXTOOLS2 function *qpDstat* with f4mode = ‘YES’ (Maier et al., 2022). Under this *f*4 configuration, perfect agreement between high and low coverage genotypes would result in *f*4 = 0, with the magnitude of the *f*4 statistic expected to increase as the low coverage genotypes become less consistent with the truth set. The direction of the *f*4 statistic can also be informative about the causal nature of the underlying discrepancies and is discussed further in the results. In these tests, the combined set of Papuan Highlander samples (38 samples) and East Asian samples (32 samples) in the global dataset were used as proxies for Papuan and East Asian ancestry, respectively, with four Mbuti samples from the SGDP project being used to represent the African ancestry (Mallick et al., 2016).

## RESULTS

### Genotype call comparisons

The mean coverage of the eight high coverage genomes ranged from ∼21x to ∼39x, with the low coverage genomes having between ∼2x to ∼4x coverage (Table S2), meaning that coverage differed by two fold within both sequencing schemes, and 10 fold across the two sequencing schemes. Merging the eight high coverage samples with the global dataset resulted in a joint variant set with 5,989,969 SNPs, with each high coverage genome having genotypes called between ∼5.27M and ∼5.53M of the total SNPs (i.e., ∼88% to ∼92% of all SNPs; Table S3). For each high coverage genome, all non-missing genotypes were used as the truth set to evaluate the performance of the four genotype calling methods on complementary low coverage datasets.

Across the four genotyping methods, the number of missing sites varied widely with the ‘naive’ genotypes having by far the most missing SNPs (≥70%), and pseudohaploid and the Impute_all method having the lowest levels of missingness (<15%) (**Figure 1**). Notably, missingness was strongly negatively correlated with coverage for naive genotypes and pseudohaploid calls, but this was considerably weaker for both imputed genotypes, reflecting the impact of imputation to recover information in regions of low coverage. This effect was most notable for the Impute_all method, emphasising the added benefit of leveraging genetic information from large cohorts of closely related individuals.

**Figure 1.**
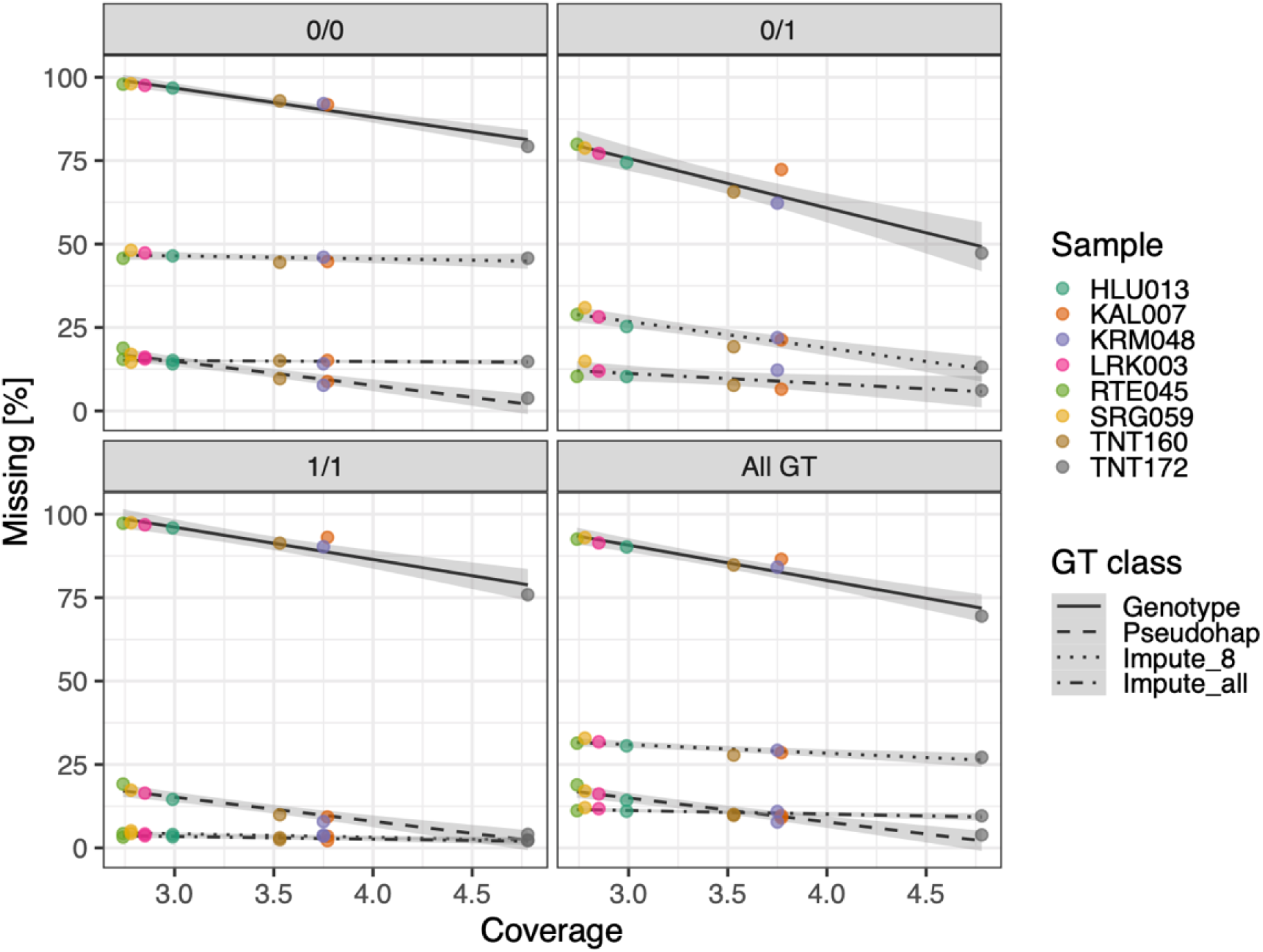
Percentage of missing sites relative to sequencing coverage for the eight low coverage samples using four different genotyping methods (see key). Results for individual and combined genotype calls are shown in panels (i.e., homozygous reference = ‘0/0’, heterozygous = ‘0/1’, homozygous alternate = ‘1/1’, and combined genotypes = ‘All GT’). Missingness is not calculable for heterozygous pseudohaploid samples, and accordingly is also not shown for combined samples.

Examination of the precision results revealed that pseudohaploids had the highest precision at homozygous calls overall (exceeding 99%; **Figure 2**), though reference bias was apparent when examining heterozygous calls (with between 0.2% to 0.4% excess of reference allele calls at these sites; **Figure 3**). Echoing the missingness results, the naive genotype calls were far less precise relative to other genotyping methods (precision between ∼55-85% across all eight samples; **Figure 2**), with the low precision being almost entirely due to large numbers of heterozygous loci that were incorrectly called as homozygous reference genotypes (Figure 2 and Table S3). Notably, both types of imputed genotypes also have their lowest precision at heterozygous sites – indicative of the general difficulty in imputing heterozygotes from low coverage data – though achieved appreciably higher precision across all three genotypes overall (>95% for Impute_8 and >98% for Impute_all across all genotypes; **Figure 2**). Precision was also positively correlated with coverage across all genotyping methods, most notably at heterozygous sites (**Figure 2**), with Impute_all again having greatly reduced dependency on coverage relative to Impute_8 genotypes, reinforcing the value of using sizable population cohorts in GLIMPSE’s imputation process.

**Figure 2.**
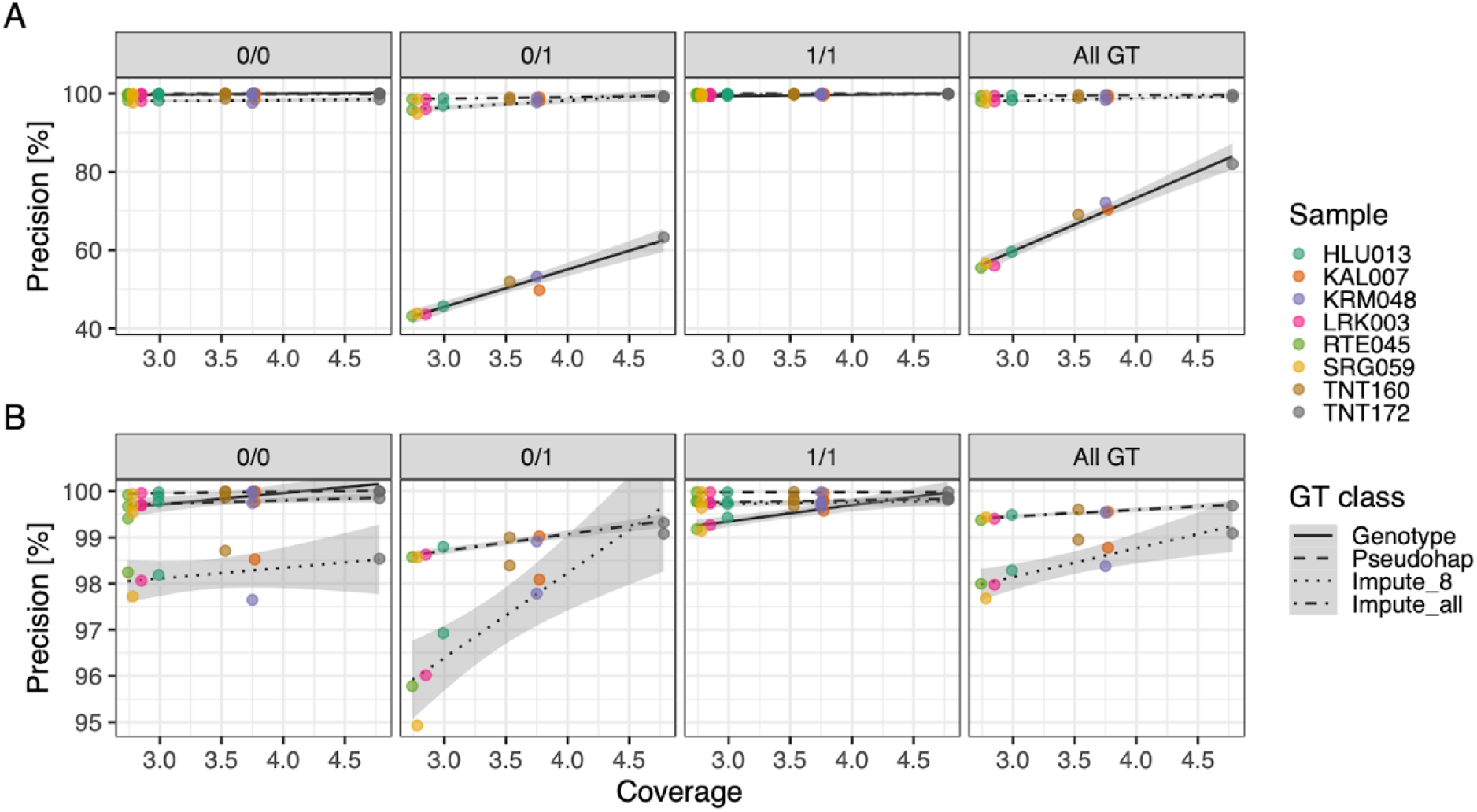
Genotyping precision relative to level of sequencing coverage for the eight low coverage all samples using four different genotyping methods. Results are shown with (A) and without (B) naive genotype calls. Precision is not calculable for heterozygous pseudohaploid samples, and accordingly is also not shown for combined samples.

**Figure 3.**
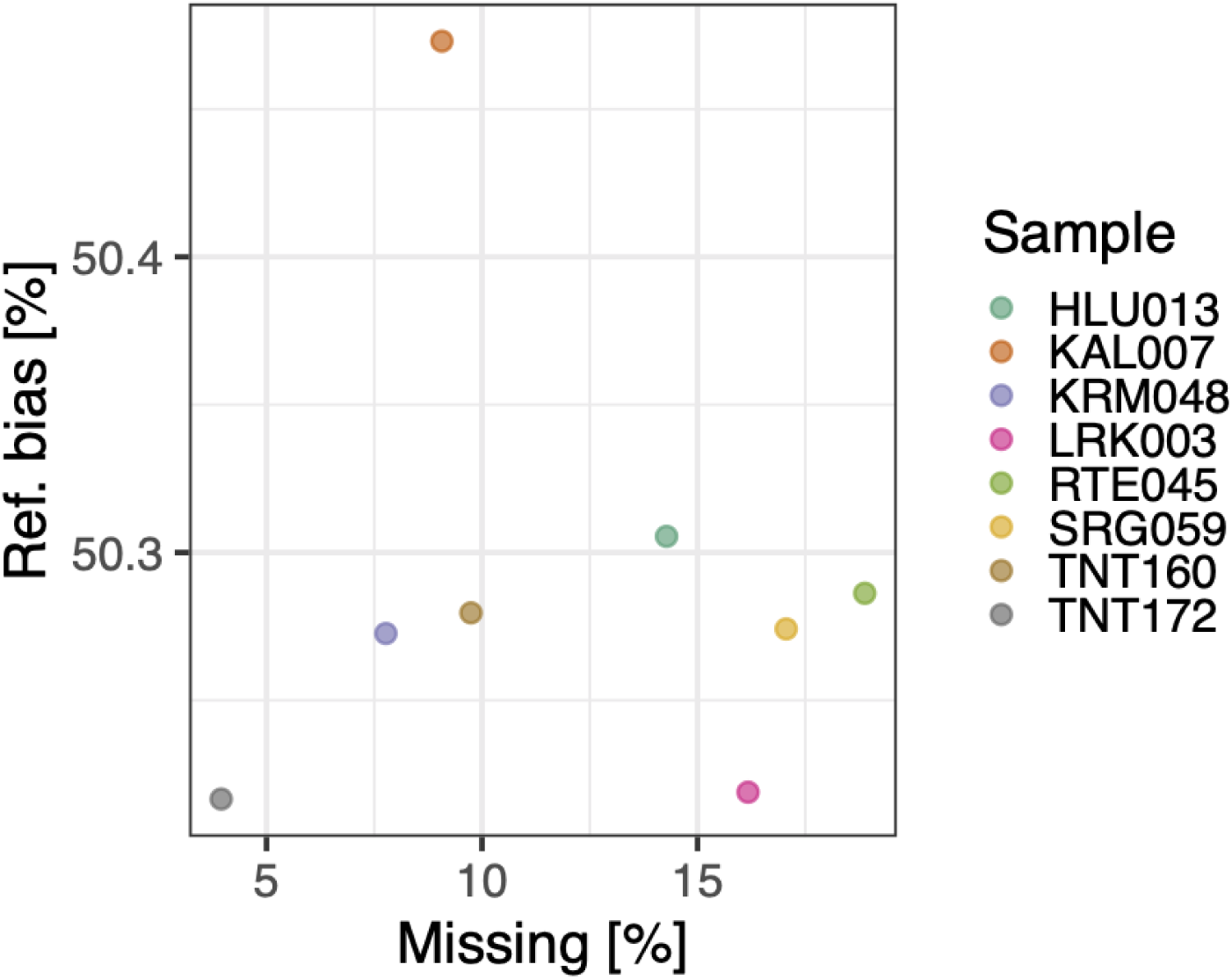
Reference bias for pseudohaploid calls from low coverage data. Reference bias measures the proportion of reference alleles sampled at heterozygous sites, which values >50% indicative of an excess reference alleles being sampled at these loci.

### PCA and ADMIXTURE

The genotype calls from low coverage genomes were projected onto principal component axes defined by the eight high coverage genomes and a global human genome dataset. This resulted in reasonable fits for pseudohaploid and both imputed genotypes, with the naive genotype calls faring the worst overall (the first four PCs are visualised in **Figure 4**). When measuring distances between genotypes from low and high coverage genomes, the Impute_all method tended to be situated closest to the truth genotypes for six of the eight samples (Figure 5), with the Impute_8 being the closest for the remaining two samples – despite the pseudohaploids being the closest match to the truth set in the first two PCs (**Figure 4**). Unlike the precision results, no correlation is evident between distance and level of sequencing coverage for the low coverage genomes, implying that coverage had negligible impact on the PCA projection of low coverage genomes in the current context.

**Figure 4.**
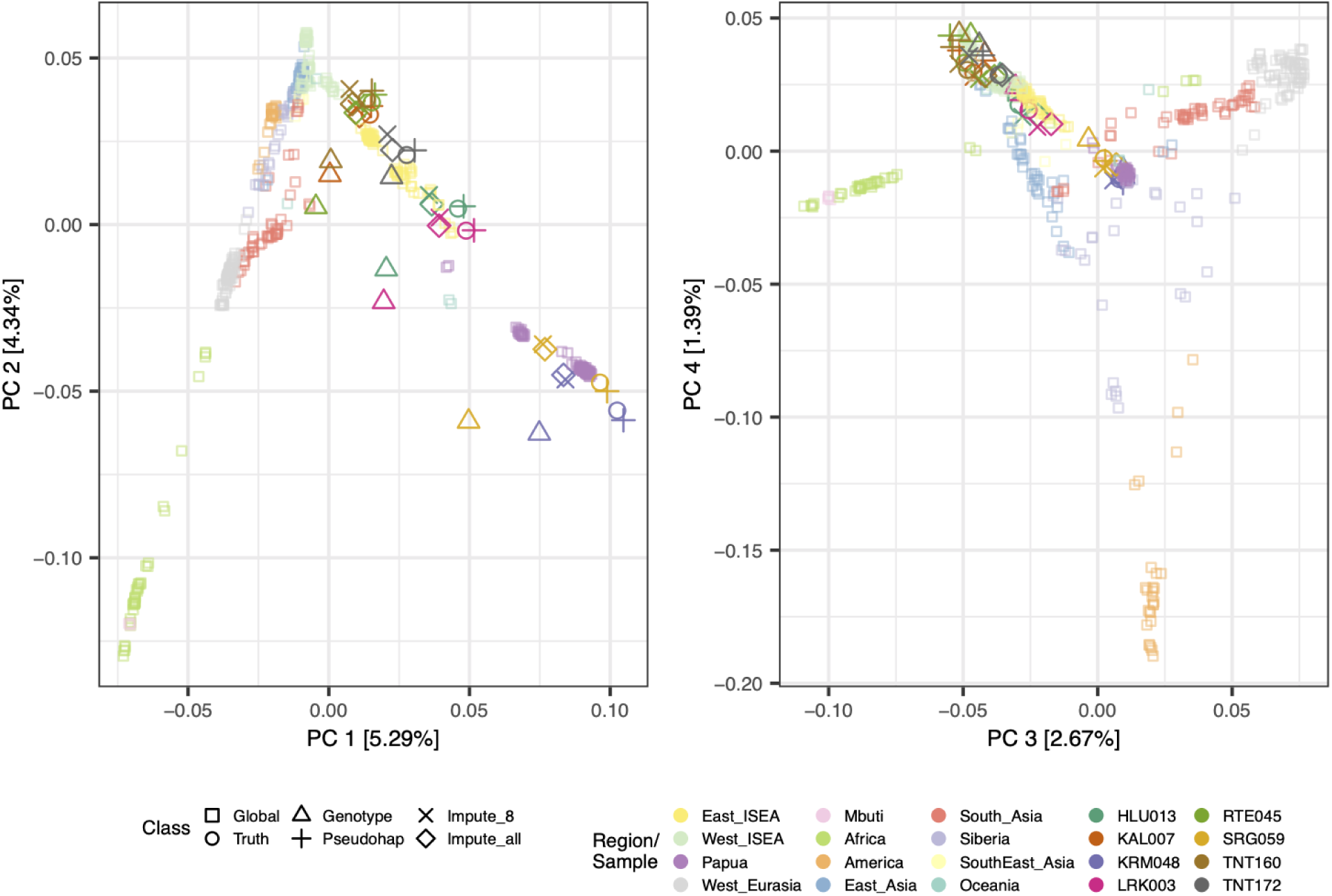
The first four principal components from a PCA based on a global human dataset and the eight high coverage Wallacean and Papuan genomes. Genotypes for low coverage Wallacean and Papuan genomes were projected onto these four PCs.

**Figure 5.**
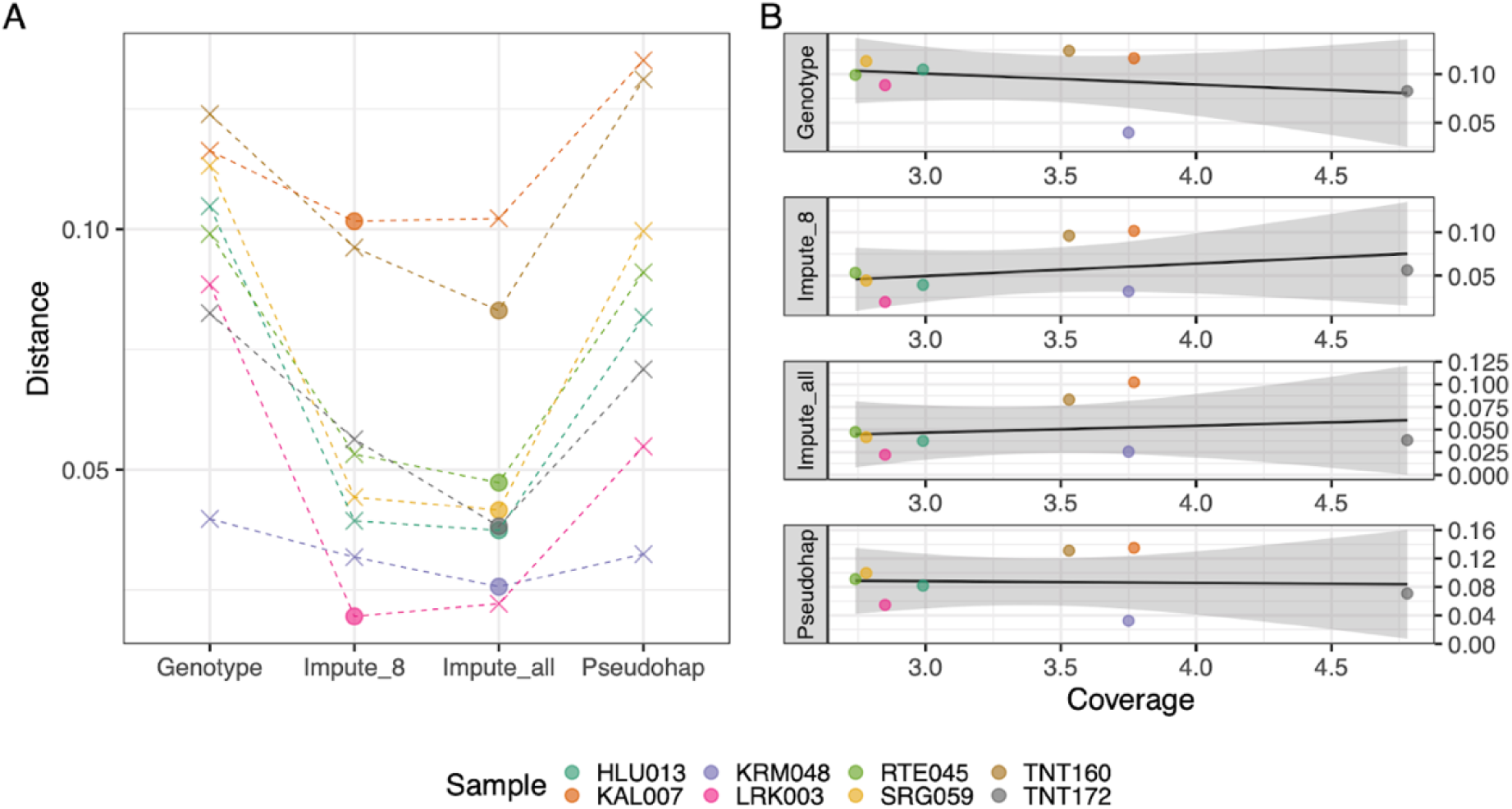
Euclidean distances between truth sets and each low coverage genotyping method calculated across PCs 1 to 10. (A) Euclidean distance was minimal (circles) for the imputation method (typically when using all ∼260 samples). (B) There were no linear associations between distance and sequence coverage.

The ADMIXTURE analyses were similar to the PCA results, in that the estimated ancestry proportions for the imputation and pseudohaploid methods provided a much better fit to the truth set than the naive call for all eight samples data (**Figures 6, 7, S1, S2**). Distance was also unrelated with sequencing coverage for the pseudohaploid and imputed genotypes; however it had a negatively linear relationship with the level of sequencing coverage for the naive genotype calls, suggesting the naive calls became proportionately worse as coverage decreased. Notably, the error in the naive genotypes tended to manifest as a deficit of Papuan and East Asian ancestry and an excess of African and South Asian ancestry relative to the truth set for all eight samples, reflecting results in the PCA where the naive genotypes are shifted away from the truth set and toward individuals with African and South Asian ancestry in the first PC (**Figure 3**). Finally, unlike the PCA results, the pseudohaploid calls provided a better fit than the two imputation methods to the true data across all samples for ancestry estimation (**Figure 7**).

**Figure 6.**
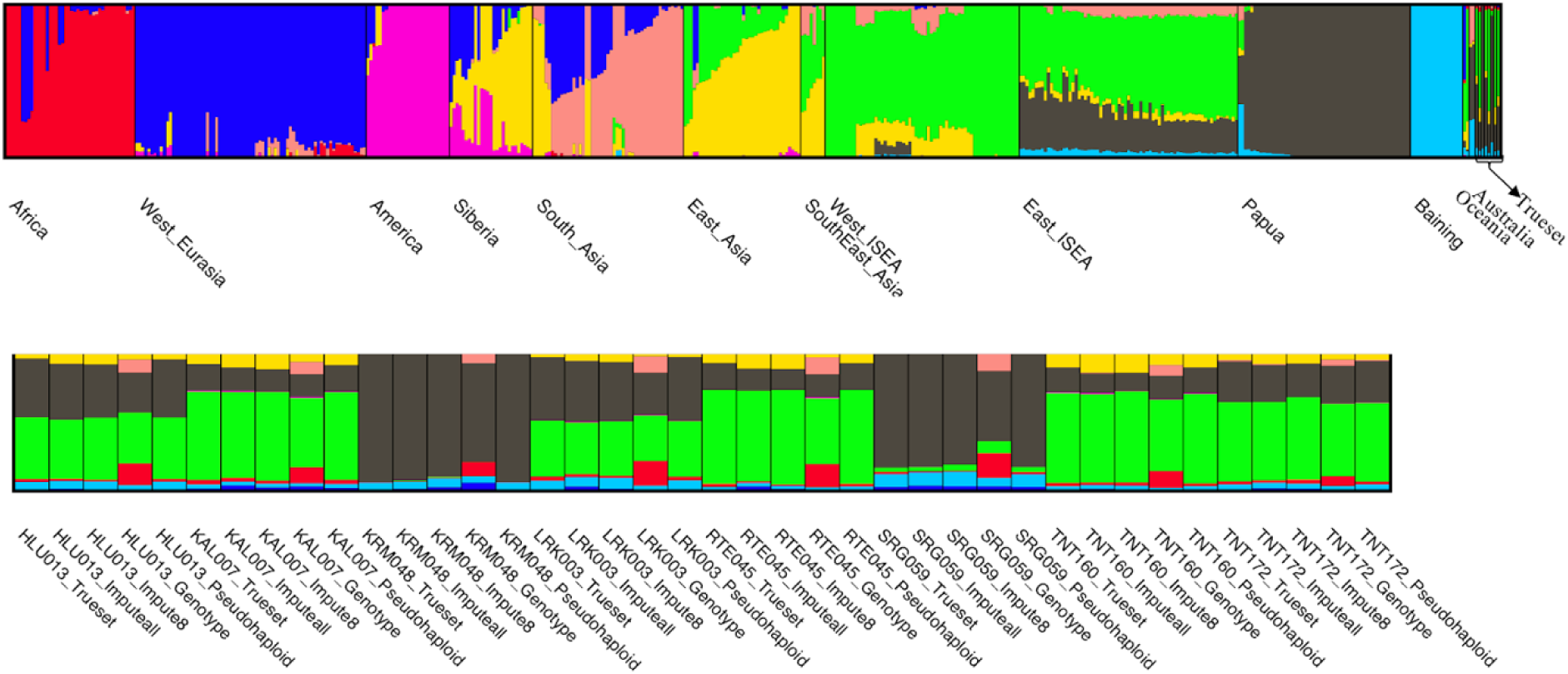
Optimal ADMIXTURE results revealed eight ancestry components amongst worldwide human samples and eight high coverage Papuan genomes (top panel). These components were used to estimate ancestry proportions for the four low coverage genotyping methods (bottom panel).

**Figure 7.**
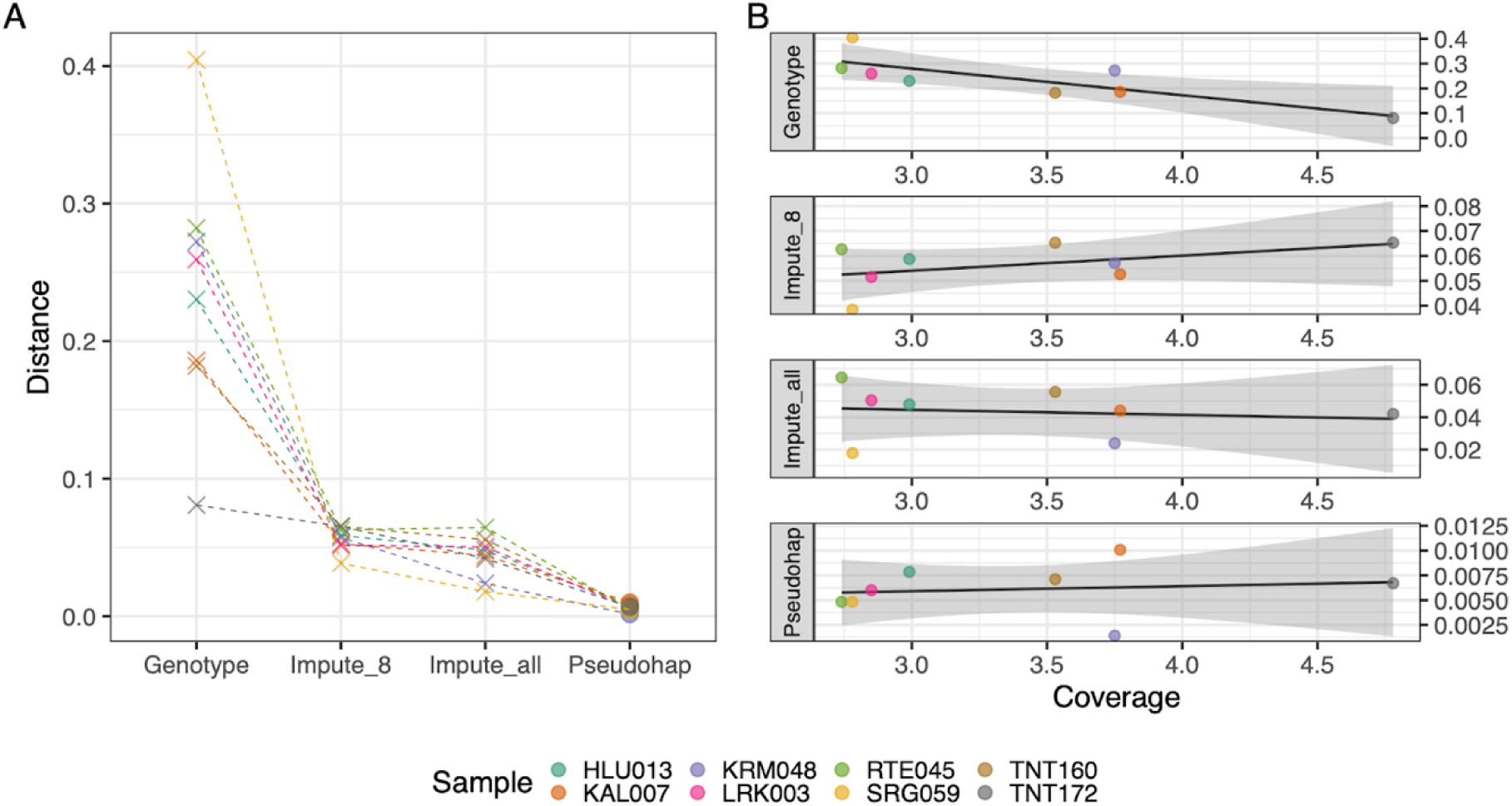
Euclidean distances between truth sets and each low coverage genotyping method calculated for optimal ADMIXTURE results (i.e., *K* = 8). (A) Euclidean distance was minimal (circles) for the pseudohaploid method. (B) A negative linear association was observed between distance and sequence coverage for the naive genotype calls, but no association was observed for other genotyping methods.

### *f*4 statistics

The *f*4 statistics reiterate the general trends observed in the PCA and ADMIXTURE analyses, with the *f*4 statistics being largest for the naive genotype calls across all eight samples, and the three other genotype methods producing *f*4 values close to 0, reflecting low and high genotype concordance with the truth genotypes, respectively (Figure 8). Significantly positive *f*4 values (absolute standardised score greater than 3 standard errors from 0; i.e., |*Z|* > 3) were observed for the naive genotypes regardless of whether the Papuan or East Asian populations were included as the Test population, with the magnitude of the *f*4 statistic for the naive genotypes tending to decrease when using the LD-pruned dataset. This result is consistent with PCA and Admixture findings – where the naive genotypes are pulled toward African samples or show excess African ancestry, respectively – which suggests that low coverage samples may appear more ‘African’ relative to the truth set as more SNPs are evaluated, resulting in larger positive *f*4 values.

**Figure 8.**
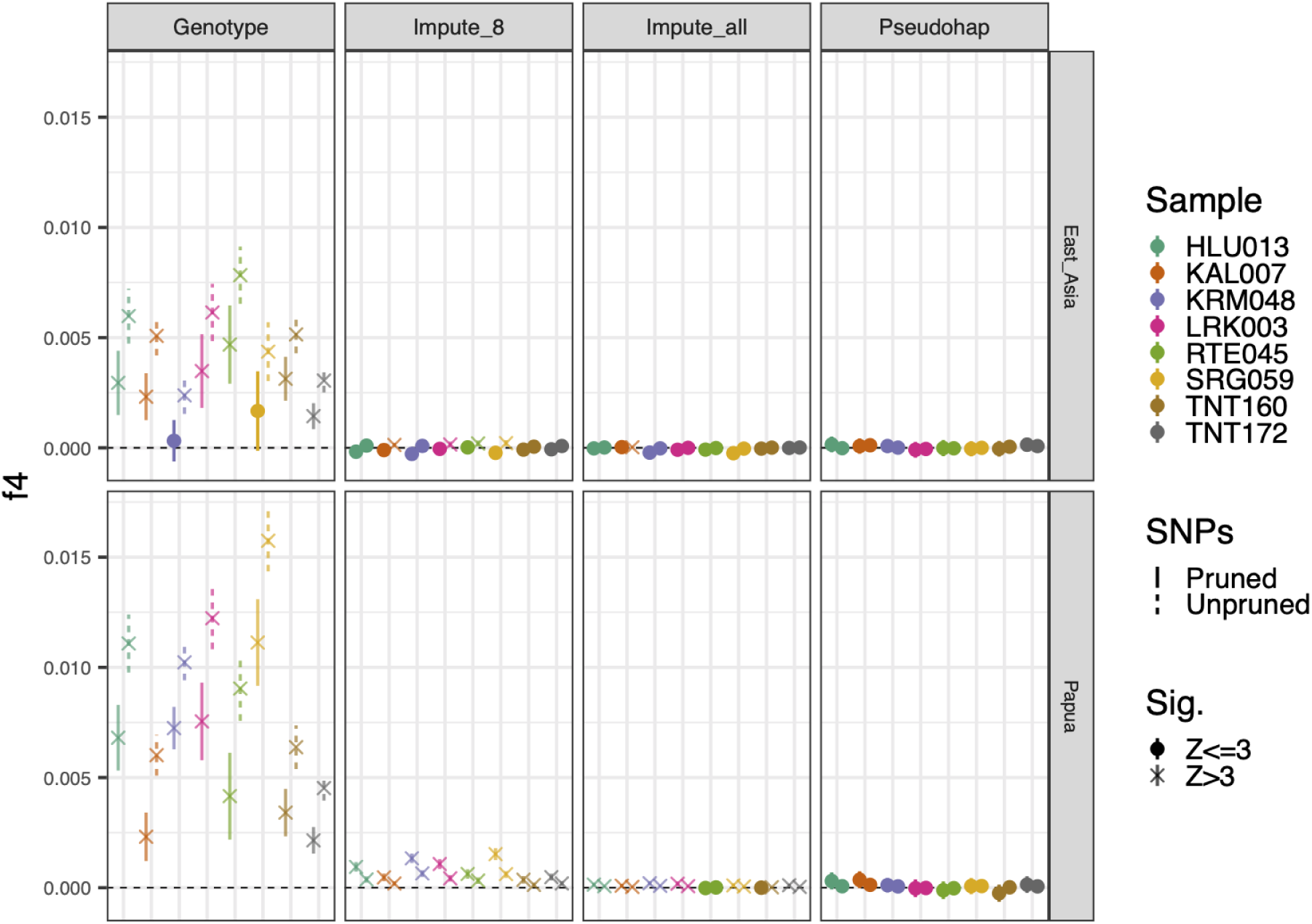
The *f*4(Truth set, Target = Low coverage; Test = Papua|East Asia, Africa) statistics calculated for four different genotyping approaches using both LD-pruned and full (i.e., unpruned) SNP sets (see key).

For the imputed genotypes, the impact of the SNP dataset on the *f*4 values depended on composition of the *f*4 population quartet. When the *f*4 statistic was evaluated using the Papuan population in the Test position, both imputed genotype methods returned positive *f4* values, but these were always lower when the full SNP set was used. When the East Asian population was used instead, *f*4 values tended to switch from positive to negative when using the unpruned or full SNP set, respectively (**Figure 9**). These results suggest that the imputation process tended to underestimate the true degree of Papuan ancestry amongst low coverage genomes but overestimated the East Asian component – possibly reflecting the lack of Papuan samples in the reference panel, such that increasing the number of SNPs either reduced or accentuated the bias in the *f*4 statistic, respectively. Echoing previous results, the Impute_all method tended to have *f*4 values much closer to zero than the Impute_8 method for all samples regardless of which SNP set was used, further emphasizing the potential for improved population genetic estimation by leveraging genetic information from population cohorts.

**Figure 9.**
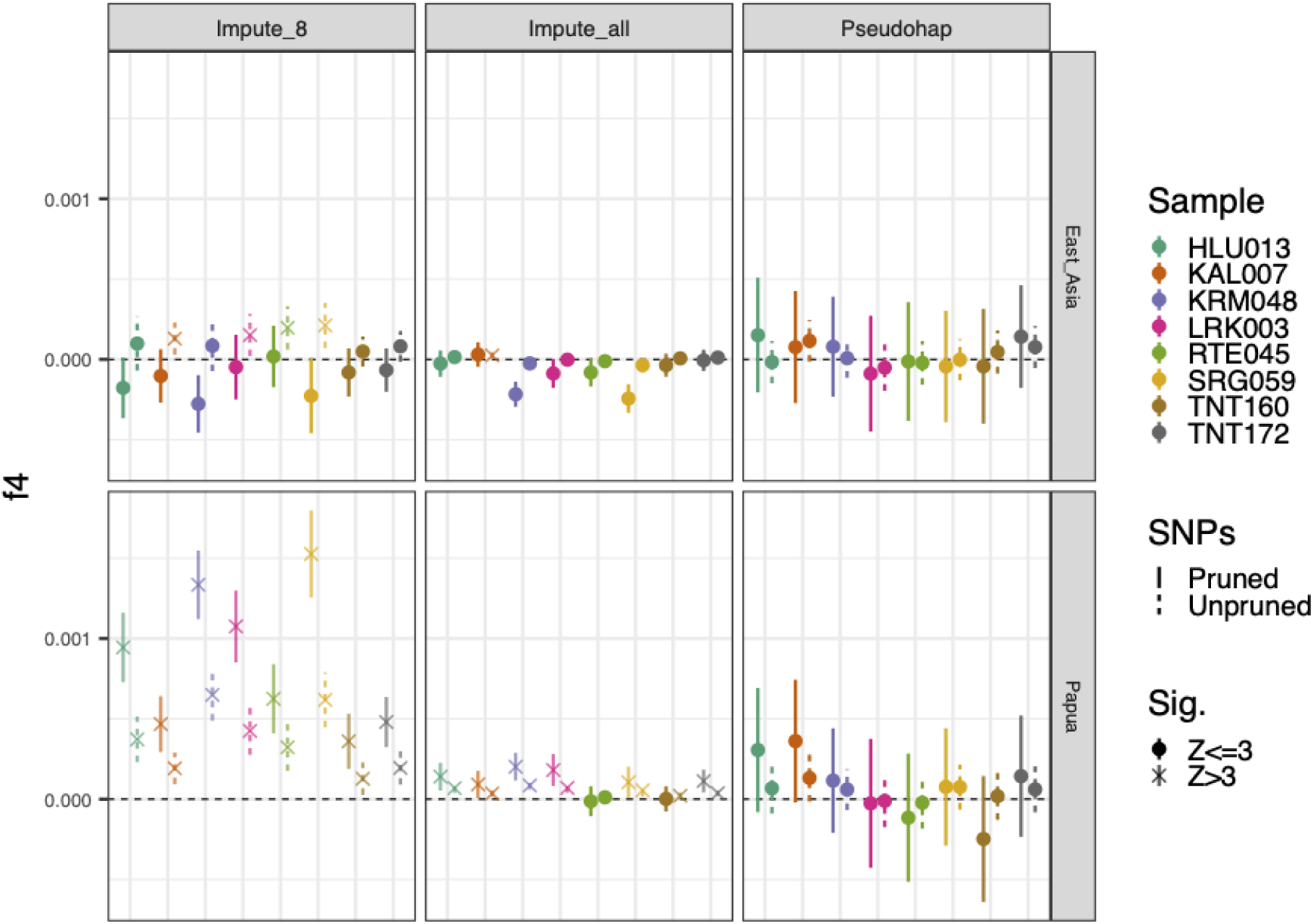
The *f*4(Truth set, Target = Low coverage; Test = Papua|East Asia Asia, Africa) statistics calculated for three different genotyping approaches (naive genotypes omitted) using both LD-pruned and full (i.e., unpruned) SNP sets (see key).

Finally, the pseudohaploid calls displayed the least bias amongst all genotyping methods, having no significant *f*4 statistics amongst all evaluated scenarios (Figure 9). While being the least biased method overall, the pseudohaploid *f*4 statistics exhibited substantially more uncertainty than the Impute_all genotypes when using comparable SNP sets. This suggests that the pseudohaploid method was more accurate, but less precise, than the imputed data for measuring *f*4 statistics under the current testing framework.

## DISCUSSION

The benchmarks demonstrate the utility of low coverage WGS in population genomic research when using pseudohaploid or imputed genotypes, achieving high precision (e.g., >98% for imputed genotypes across ∼6 million SNPs) and robust inferences for PCA, ADMIXTURE and *f*4 analyses that align closely to the truth sets. In general, inferences from imputed genotypes, particularly those leveraging genetic information from cohorts of related individuals in the target populations (i.e., Impute_all), were comparable to pseudohaploid calls – which are generally regarded to be highly robust to statistical artefacts resulting from SNP ascertainment (Lachance & Tishkoff, 2013) – with the best choice differing depending on the statistic. Importantly, the results of the *f*4 tests in this study suggest improved accuracy of pseudohaploid calls over the imputed genotypes observed for some statistics (i.e., first few PC dimensions, ADMIXTURE, *f*4) may ultimately come at the cost of lower precision (**Figure 9**). In other words, while pseudohaploids produce results that are closer to the expected value on average than the imputed genotypes, they also exhibit more variation, meaning that a single estimate from pseudohaploid data will frequently be further from the truth than an estimate made from imputed genotype data. This tradeoff between precision and accuracy is a fundamental property of statistical estimation, with lowered precision of pseudohaploid data likely stemming from only having half the information of standard biallelic genotypes and the imputed genotypes prone to bias that is dependent upon the composition of the reference data.

Importantly, recent analyses of GLIMPSE imputation on simulated genomic datasets (Lou et al., 2021) indicate that more precise genotype imputation should be possible from what is reported here, simply by increasing the number of samples in the target population. In the current study, the number of samples per target population is relatively small (∼20 to ∼30), so it was not possible to examine potential gains from having 100s to 1000s of samples (or more). In particular, it is important to understand the trade-off between sequencing coverage and sample size in target populations when using GLIMPSE – i.e., whether to sequence a handful of samples at reasonably high coverage vs sequencing more samples at proportionately lower coverage. This question has been examined for other low coverage WGS imputation algorithms (e.g., STITCH; (Davies et al., 2016)) – but, to our knowledge, this question has not been explored when the imputation algorithm also leverages information from a genomic reference panel. Similarly, it remains to be seen how much GLIMPSE imputation performance can be improved by having more informative reference panels, and how these gains affect, and are affected by, target population size. For instance, the recently published Genome Asia Panel (GenomeAsia100K Consortium, 2019) provides a more representative dataset than the TGP for the current study, however these genomes were not phased at the time of this study so were not suitable for the GLIMPSE imputation process.

Finally, while we have only evaluated ‘hard’ genotype calls here, recent studies have emphasised the utility of using genotype likelihoods (GL) or dosages in the place of hard calls (i.e., assuming a single genotype) for population genomic inference. Many GL-based population genomic inference procedures now exist (e.g., ANGSD, (Korneliussen et al., 2014)) and have the advantage of incorporating underlying uncertainty in the genotype calls into downstream estimates. We are not aware of any published research that explores the performance of GL inference on imputed genotype datasets, though this could result in further improvements in inference by appropriate quantification of uncertainty. Indeed, this may be partly because the adoption of GL-based inference methods has been slow across the population genomic community in general, possibly because of the relative novelty of the methods and the lack of suitable published GL datasets.

### Conclusion

Studies of increasingly larger datasets are continuing to emerge as sequencing costs steadily decrease, though standard laboratory budgets mean that SNP panels have remained the most common way to produce population genomic data for cohorts exceeding dozens of individuals. Our study highlights the promise of low coverage sequencing in combination with genotype imputation to facilitate robust population genomic analyses for human groups that lack suitable reference datasets and suitable SNP panels. Ongoing improvements to the GLIMPSE algorithm have led to increased power to accurately capture rare alleles when using suitable reference datasets with 100,000’s of genomes (Rubinacci et al., 2023), though our results show that GLIMPSE can produce robust population genomic inferences even when reference panels are not representative of the target population when genetic information from the targeted individuals is used. The increasing power and broad utility of this approach means it will likely be increasingly used in place of SNP panels for statistical and population genomic research in the future.

---

**Figure S1.**
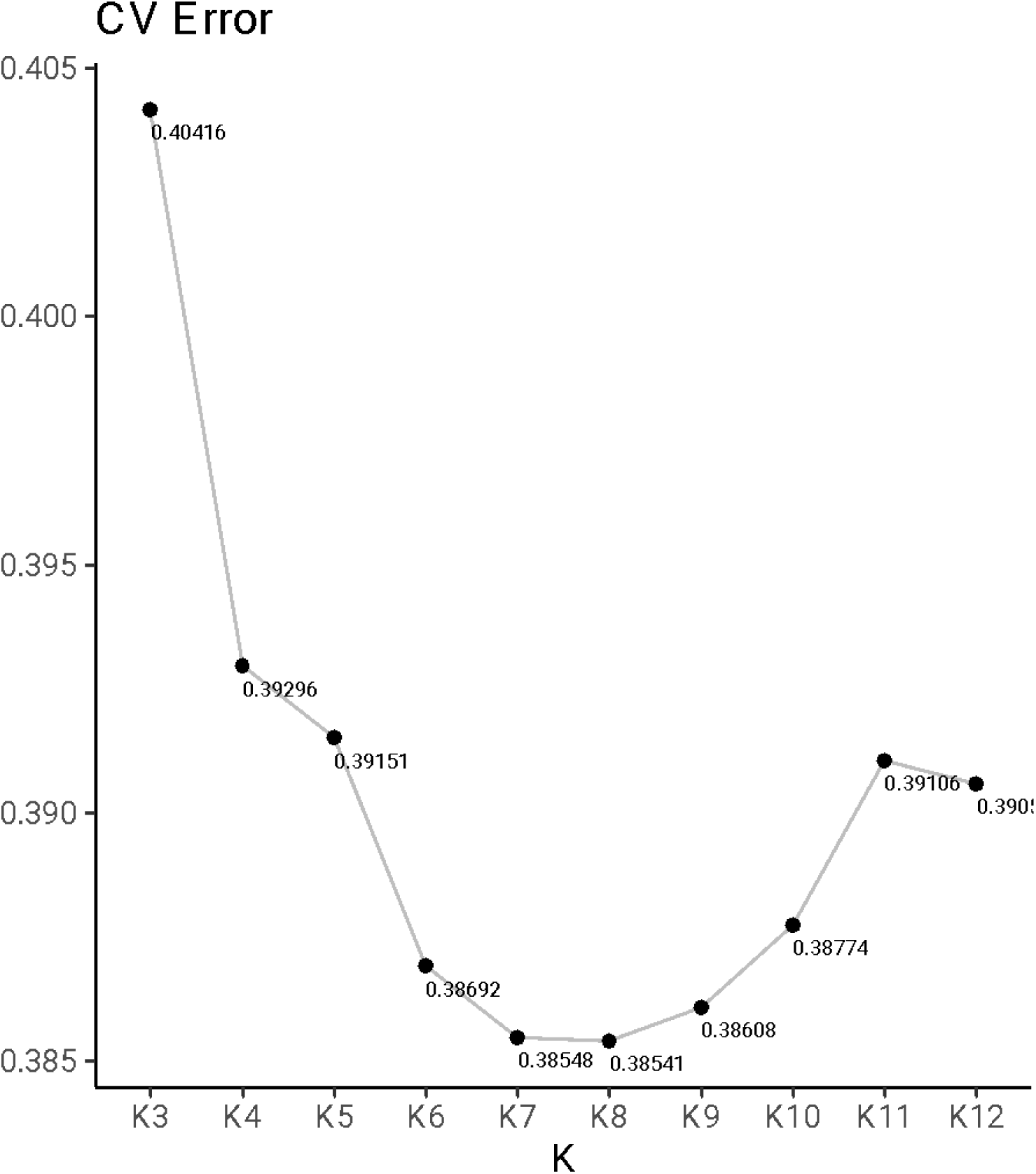
Cross-validation error for ADMIXTURE analyses of the global dataset. The optimal fit was *K* = 8 components, with testing performed for *K*=3 to *K*=12 components.

**Figure S2.**
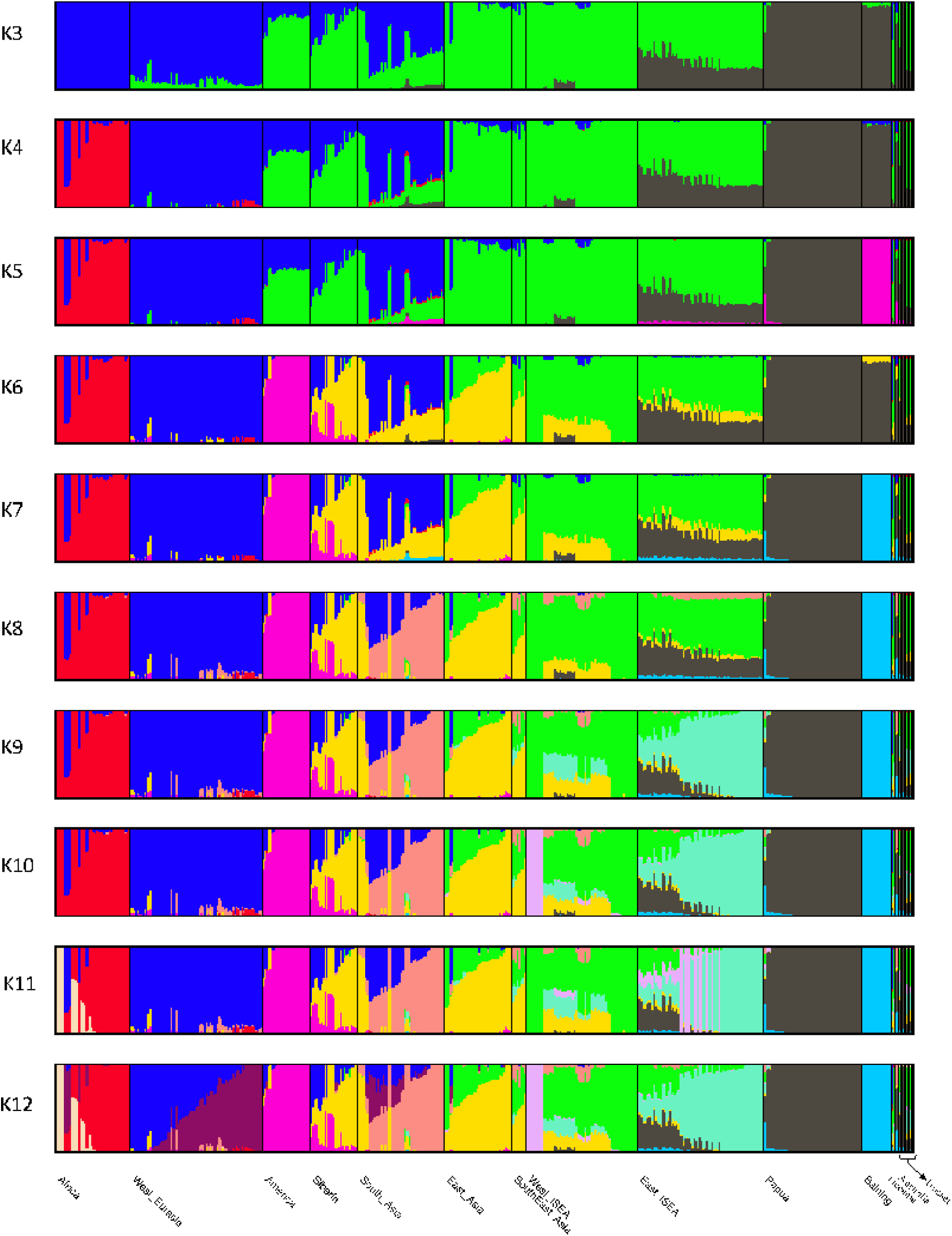
Ancestry composition for individuals in the global dataset for 3 ≤ *K* ≤ 12 distinct components. Model fit was determined using the minimum cross-validation error (see Figure S1).

